# Repurposing of an Antifungal Drug against Gastrointestinal Stromal Tumors

**DOI:** 10.1101/2021.01.15.426618

**Authors:** Charli Deepak Arulanandam, R Prathiviraj, Govindarajan Rasiravathanahalli Kaveriyappan

## Abstract

Drug discovery is an important research area to improve human health. Currently, treatment of gastrointestinal stromal tumors (GISTs) is unsuccessful due to drug-resistance, hence, there is a demand for alternatives. Often, there is limited time available for toxicological assessments and a lack of safer drugs. It is possible to identify new drugs from existing approved drugs possessing another purpose in the clinical lines. In this study, virtual screening of some Food and Drug Administration (FDA-USA) approved and available antifungal and antineoplastic drugs were performed against GISTs based on docking affinity of human platelet-derived growth factor receptor alpha (PDGFRA) with these drugs to identify a suitable PDGFRA inhibitor for saving the time required for toxicity screening. The protein and ligand-binding affinity were investigated for five FDA approved antineoplastic and thirty-six antifungal drugs against PDGFRA using the AutoDock (AD) and AutoDock Vina (ADV) software. Based on docking score and inhibition constant (K_i_), Itraconazole was predicted as a better PDGFRA inhibitor among all the computationally tested drugs.

## INTRODUCTION

Gastrointestinal stromal tumors (GISTs) are rare primary neoplasms of the gastrointestinal tract, mesentery, or omentum. They originate from the interstitial cells of Cajal (ICC) or their progenitor cells and are the most common mesenchymal neoplasms in the human digestive tract [1][2]. Platelet-derived growth factor receptor alpha (PDGFRA) is part of a family of proteins called receptor tyrosine kinases (RTKs). GISTs cause mutation in Exon 18 which can activate D842V from aspartic acid (D) to a valine (V) by an amino acid substitution at position 842 in the PDGFRA [3]. It is a very rare subgroup of GIST (about 10 %) known to be resistant to conventional tyrosine kinase inhibitors (TKIs) and shows an indolent behaviour [4]. Somatic mutations acquired during a person’s lifetime leading to sporadic GISTs are present only in tumor cells. The annual new incidences were 10,660 and death toll was estimated to be 3820 for 2009 in the United States, including adults and children [5]. In Caucasian populations, their annual GISTs are 10 to 15 cases per million inhabitants. However, GISTs may be more common than this estimate suggests because small tumors may remain undiagnosed. PDGFRA gene mutations associated with GISTs create a protein that no longer requires the binding of the platelet-derived growth factor protein to be activated [6]. As a result, PDGFRA protein and the signalling pathways are constitutively activated which increases cell proliferation and survival, leading to the formation of a tumor. The World Health Organization (WHO) classification of tumors of lymphoid tissues and hematopoietic cells introduced a new category for myeloid, lymphoid neoplasms with eosinophilia and abnormalities of PDGFRA [7]. As in GISTs, the constitutively active PDGFRA protein leads to an overgrowth of cells and the formation of tumors. PDGFRA (wild-type GISTs) generally shows primary resistance to Imatinib [7]. Although the marketed pharmaceutical Imatinib is thought to be the most effective agent for treating GISTs, about half of patients with unresectable or metastatic GISTs develop secondary resistance within two years of Imatinib therapy. Most patients develop resistance or intolerance to Regorafenib within a year [8]. Drug-resistance and failure to treat GISTs tumors, therefore, warrant the need to develop new inhibitors for GIST treatment. Selection of antifungal agents as alternative drugs for GISTs among the different FDA-approved antifungals using *in silico* analysis is a promising, time effective and cost-saving approach for repurposing already marketed drugs. Their pharmacokinetics and other properties were already approved by regulatory bodies for new purposes [9]. Thus, therapeutic switching is a promising approach to provide anticancer drugs. Drugs developed for other purposes such as Mebendazole (antihelminthic), Nitroglycerin (vasodilator), Cimetidine (H_2_ receptor antagonist), Clarithromycin (antibiotic), Diclofenac (a non-steroidal anti-inflammatory drug), and Itraconazole (antifungal) were identified for their anticancer properties. The antifungal agent Ciclopiroxolamine (CPX) demonstrated promising antineoplastic activities against haematological and solid tumors [10]. PDGFRA protein signalling is important for the development and regulation of cell proliferation and cell survival throughout the body. Studies suggest that PDGFRA plays an important role in organ development, wound healing, and tumor progression. Mutations in its gene have been associated with the idiopathic hypereosinophilic syndrome, somatic and familial gastrointestinal stromal tumors, and a variety of other cancers (https://www.proteinatlas.org/ENSG00000134853). Depending on the context, these drugs promoted or inhibited cell proliferation and cell migration (https://www.uniprot.org/uniprot/P16234).

Molecular docking approaches explore the receptor-ligand conformations within the binding sites of macromolecular targets. Structure-based drug discovery is widely used by the scientific community in Medicinal Chemistry to estimate the ligand-receptor binding free energy by evaluating critical phenomena involved in the intermolecular recognition process. A variety of docking algorithms are currently available [11].

Thus, the objective of the present study was to evaluate FDA approved drugs for cancer treatment using *in-silico* analysis. Herein, the concept of therapeutic switching was explored using AutoDock (AD), and AutoDock Vina (ADV) docking programs to find an alternative drug for GISTs among the different FDA-approved antifungal and antineoplastic drugs.

## MATERIALS AND METHODS

### Molecular Docking

The virtual molecular screening was used to dock small macromolecules in order to discover lead compound. This *in silico* screening is an already established approach in computer-aided drug discovery [12]. A docking experiment with 36 antifungal drugs and 5 antineoplastic drugs was made by screening the protein (5K5X) and ligand interactions using AD and ADV [12].

### Cygwin

Cygwin was used to perform AD and ADV molecular docking in the Windows operating system [13]. Cygwin Packages are publicly available for docking operations at https://www.cygwin.com.

### *AutoDock (AD) and AutoDock Vina* (ADV)

AD 4.2 suite is a free software for virtual screening and computational docking. This suite includes several complementary tools [14]. ADV provides a straightforward scoring function and rapid gradient-optimization in the conformational search. ADV tends to be faster than AutoDock 4 by orders of magnitude [15]. In the docking experiment, the grid box dimension values were adjusted up to dimensions (in □ngstrØm) Grid center X:68.8375, Y:37.8355, Z:3.108 Grid Size X:33.7535, Y:33.7535, Z:33.7535 with an exhaustiveness value=8. All the compounds were docked separately against the crystal structure of human PGDFRA and the generated docked complexes were evaluated on the basis of lowest binding energy (kcal/mol) values and hydrogen interaction patterns visualized by the BIOVIA Discovery Studio (DS) [16]. The 2D graphical representation of screened compounds was obtained from BIOVIA DS based on the best docking score/binding energy from the tested antifungal and antineoplastic drugs.

### Preparation of target protein

Human PDGFRA is built of a single subunit (Figure 1). The crystal structure of human PDGFRA was acquired from Protein Data Bank as PDB format (PDB ID: 5K5X - *Homo sapiens*) with PDGFRA classified as a transferase in the RCSB PDB database accessible at web server https://www.rcsb.org/. Before preparing the protein PDBQT format in AutoDock, the structure of the target protein was validated using the software MolProbity [17]. This server is accessible from http://molprobity.biochem.duke.edu. Based on this server, the protein structure was solved by X-ray diffraction at 2.168 Å resolution with R-value work = 0.192; R-value free = 0.224. It has one unique chain (chain A) and in total 345 residues and 151 hetero-groups.

**Fig. 1.**
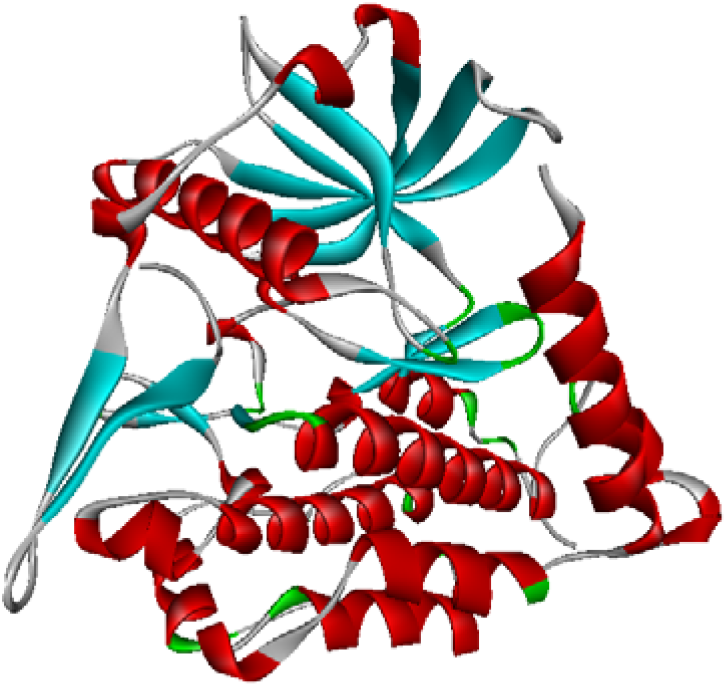
Human PDGFRA (PDB: 5K5X) structure visualized by BIOVIA DS.

### MarvinSketch

MarvinSketch was used to generate the 3D structures of the test compounds from the chemical drawing (https://chemaxon.com).

### Ligands for docking study

All the ligands and their 3D structures were shown in Fig. 2 and 3. Interactions between receptors and ligands were analyzed by AD and ADV [15]. Results of docking and bonding interactions were analyzed by BIOVIA DS [16].

**Fig. 2.**
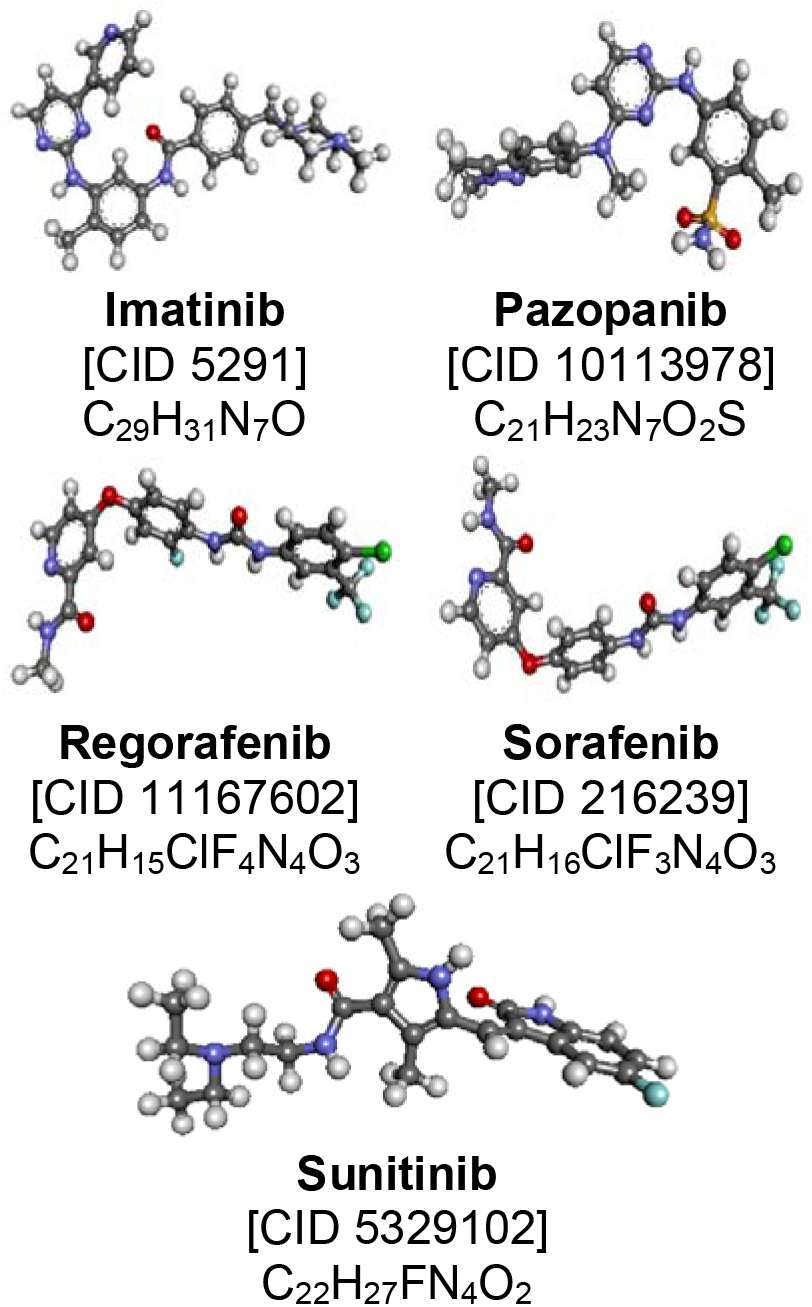
3D Structure of approved antineoplastic drugs with PubChem Identifier.

**Fig. 3.**
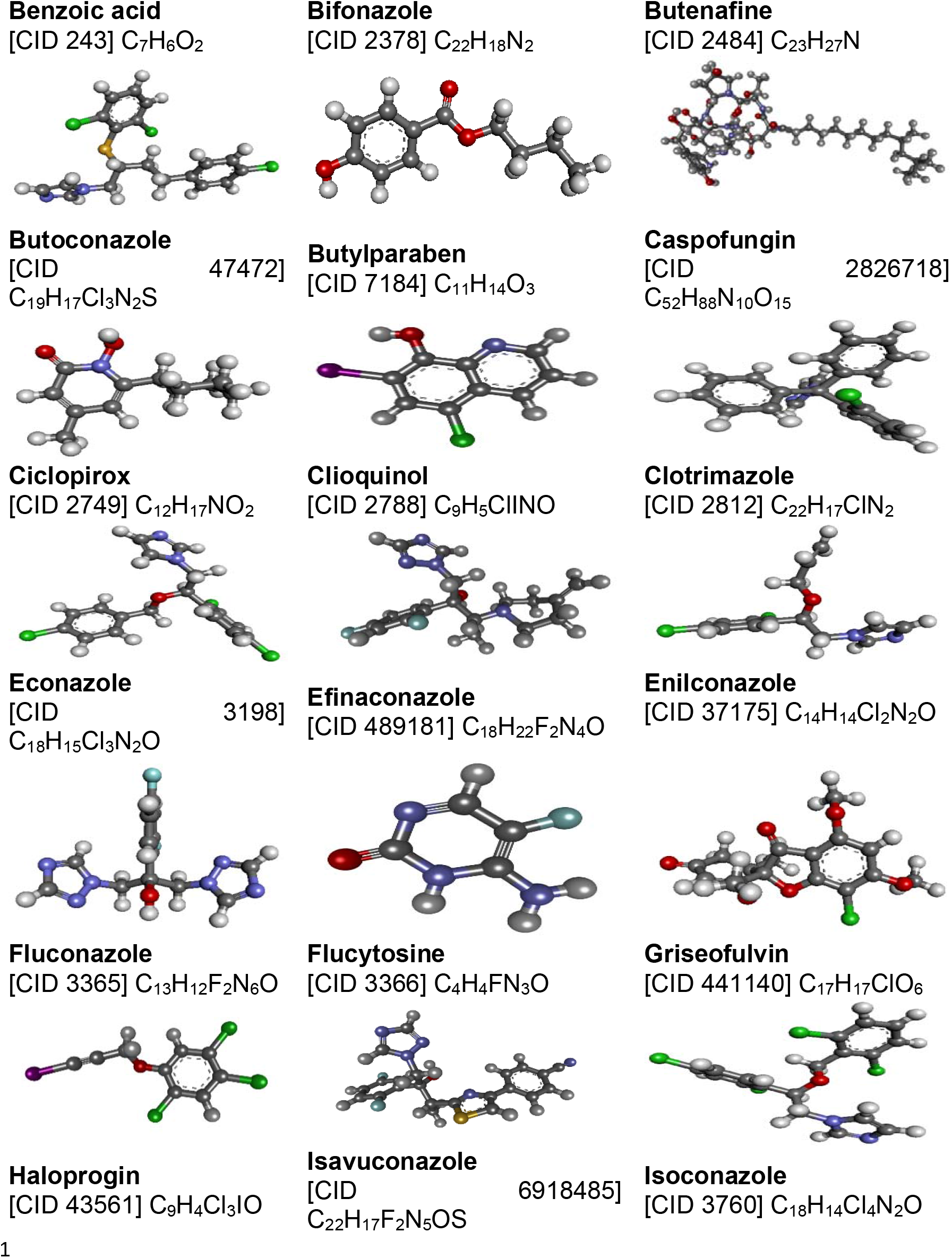
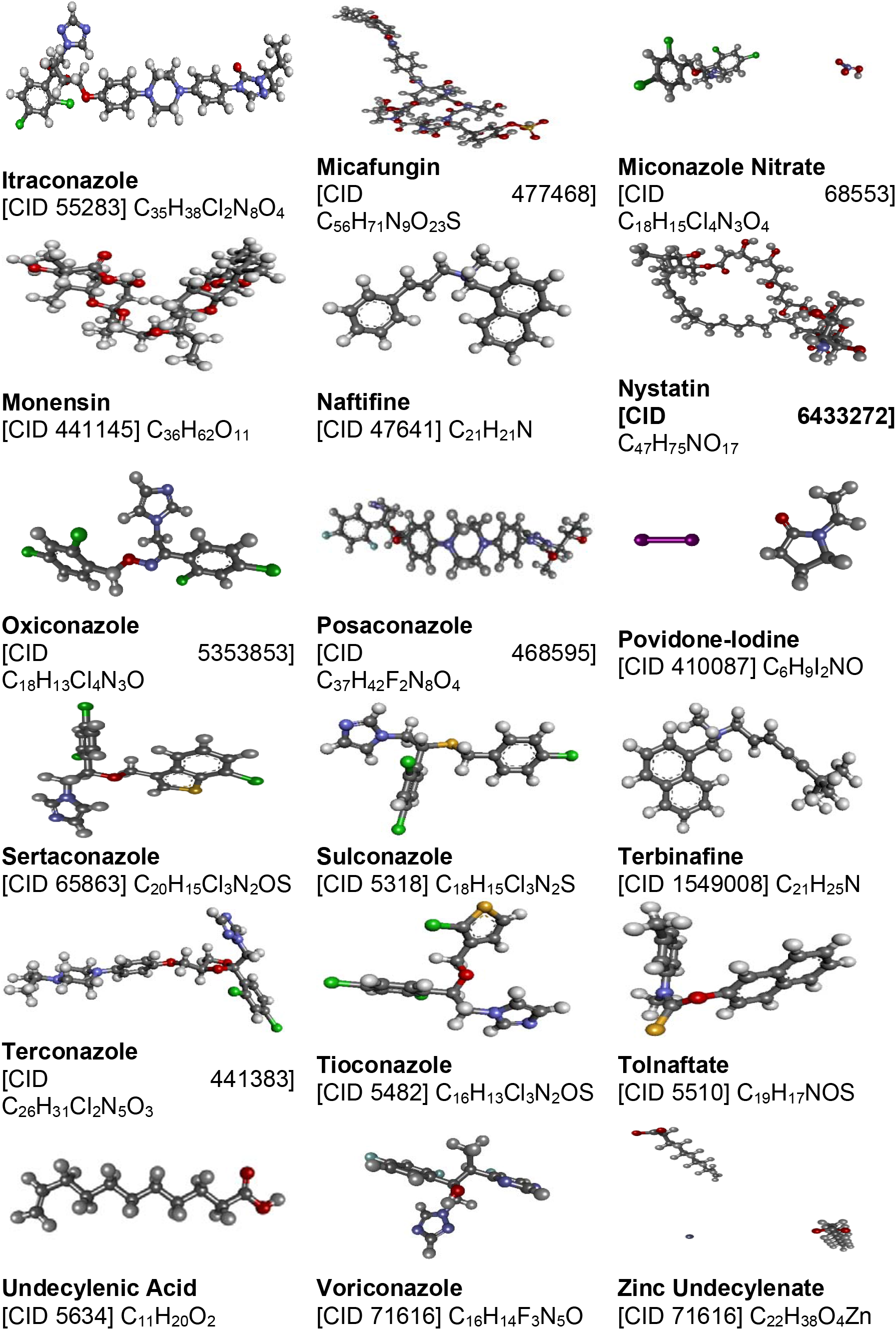
3D Structure of approved antifungal and antineoplastic drugs with PubChem Identifier.

### Antineoplastic drugs

Five antineoplastic compounds were obtained from the PubChem database. All the tested compounds were downloaded in the 3D format as SDF files. BIOVIA DS was used to visualize and convert 3D SDF files into the PDB file format for AD Vina compatibility as shown in Fig. 2.

### Antifungal drug

Currently, the therapeutic switching of existing antifungal drugs used for anti-cancer applications could be used to demarcate alternative drug potentials for the improvement of cancer therapy. Non-cancer drugs demonstrated anti-cancer activity in preclinical trials [18]. In the present work,

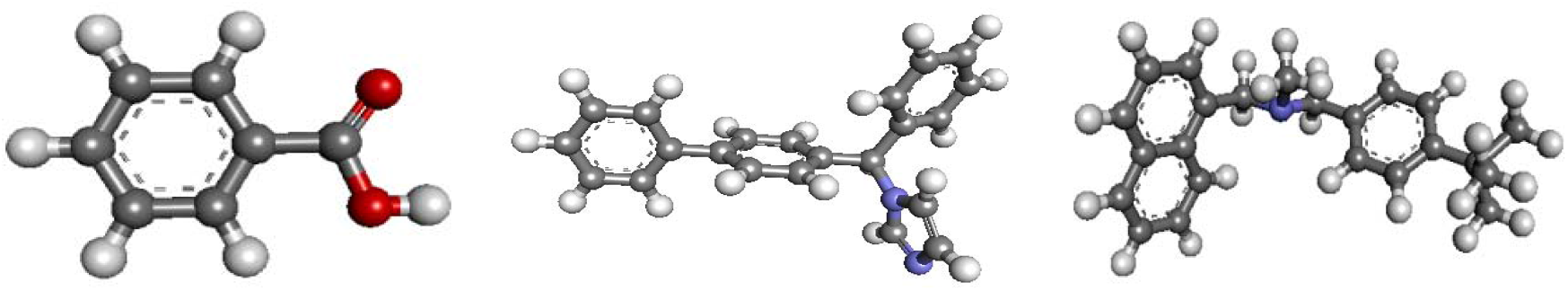

36 antifungal drugs were studied for their potential binding affinity towards human PDGFRA. As shown in Fig. 3, the 3D structure of these antifungal drugs was obtained from the PubChem database. BIOVIA DS was used to convert SDF files into PDB format.

### Ligand interaction diagrams

2D structure of the protein-ligand interaction was visualized using BIOVIA DS to know the position of the ligand bonding with amino acids present in the target protein.

## RESULTS

Human PDGFRA and all the target compounds were docked using AD, and ADV. According to AD prediction, Itraconazole docked with human PDGFRA showed the best binding energy of −7.78 kcal/mol among all tested antifungal compounds (Fig. 4). Its estimated inhibition constant (Ki) is 1.98 μM (micromolar). This inhibition constant and binding energy are better than the Sunitinib binding energy −7.34 kcal/mol with PDGFRA and Sunitinib inhibition constant was 4.14 μM. Sunitinib can be used to treat patients with Imatinib-resistant GISTs [19]. From the AD prediction, Imatinib showed a very low inhibition constant of 65.97 nM.

**Fig. 4.**
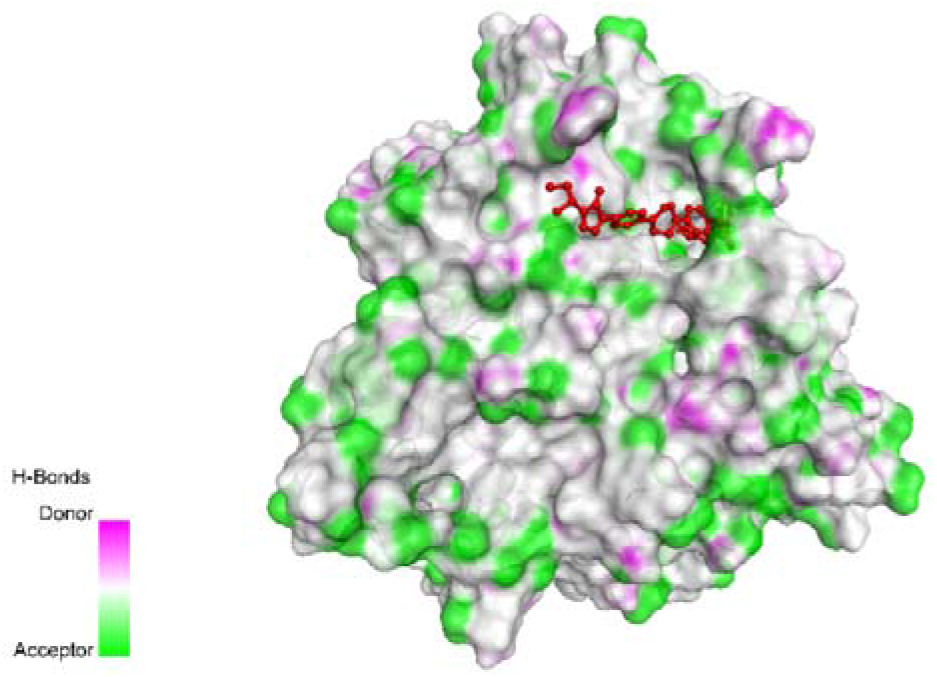
General view of PDGFRA and Itraconazole binding pose (ligand represented as a red-colored ball-and-stick model).

The 2D structure of protein-ligand interaction was obtained by applying BIOVIA DS. The protein-ligand interaction for Itraconazole is shown in Fig. 5.

**Fig. 5.**
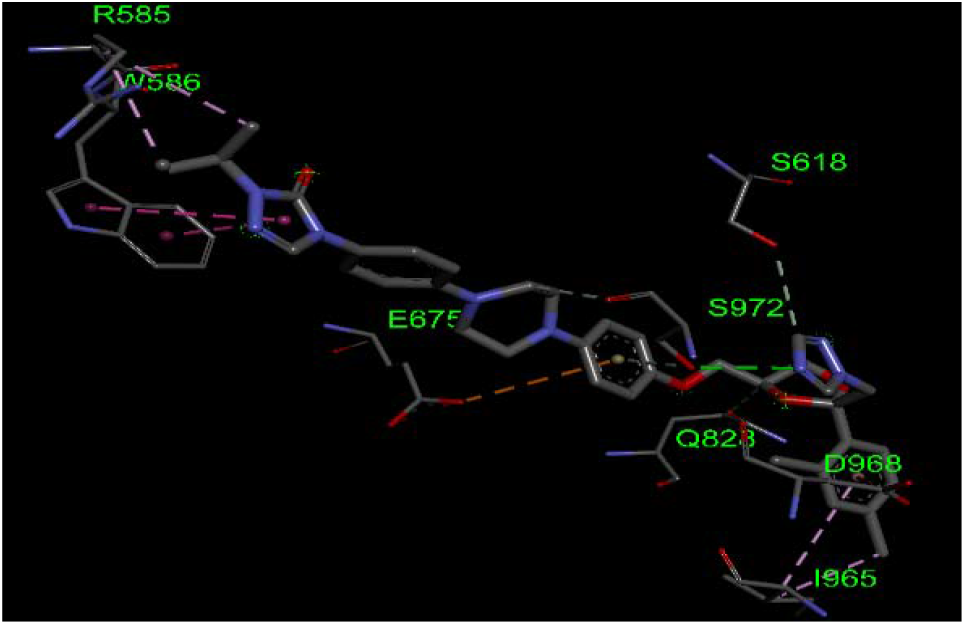
Detailed view of a docking site for the screened Itraconazole against human PDGFRA.

According to ADV docking results, Itraconazole (−9.6 kcal/mol) showed lesser binding energy than Imatinib (−9.4 kcal/mol). This was the lowest binding energy among all the test compounds. Collectively, these results suggest that Itraconazole may provide a promising inhibitor of PDGFRA. The protein and ligand binding energy for all the tested compounds were summarized in Table 1.

**Table 1.** Binding affinity of ligands and drugs

| **Ligand** | **AD binding affinity** |  | **ADV kcal/mol** |
| --- | --- | --- | --- |
|  | **kcal/mol** | **K<sub>i</sub>** |  |
| **Antifungal drugs** |  |  |  |
| Benzoic acid | -5.09 | 185.41 μM | -4.7 |
| Bifonazole | -7.56 | 2.89 μM | -8.2 |
| Butenafine | -7.42 | 3.66 μM | -8.0 |
| Butoconazole | -6.17 | 29.87 μM | -6.3 |
| Butylparaben | -5.56 | 84.49 μM | -4.8 |
| Caspofungin | -0.53 | 410.66 mM | -1.3 |
| Ciclopirox | -6.62 | 13.97 μM | -6.3 |
| Clioquinol | -6.75 | 11.31 μM | -5.4 |
| Clotrimazole | -6.46 | 18.49 μM | -6.8 |
| Econazole | -6.85 | 9.51 μM | -7.6 |
| Efinaconazole | -5.82 | 53.93 μM | -7.0 |
| Enilconazole | -5.70 | 66.50 μM | -5.9 |
| Fluconazole | -4.68 | 368.89 μM | -6.5 |
| Flucytosine | -4.68 | 372.33 μM | -4.8 |
| Griseofulvin | -6.64 | 13.61 μM | -6.6 |
| Haloprogin | -6.27 | 25.46 μM | -4.8 |
| Isavuconazole | -7.73 | 2.17 μM | -8.5 |
| Isoconazole | -7.44 | 3.51 μM | -7.2 |
| **Itraconazole** | **-7.78** | **1.98 μM** | **-9.6** |
| Micafungin | -2.65 | 11.34 μM | 27.8 |
| Miconazole Nitrate | -2.70 | 10.42 μM | -3.8 |
| Monensin | -5.14 | 170.30 μM | -8.7 |
| Naftifine | -7.92 | 1.56 μM | -7.8 |
| Nystatin | -6.00 | 40.15 μM | -7.2 |
| Oxiconazole | -7.64 | 2.52 μM | -7.4 |
| Posaconazole | -6.22 | 27.38 μM | -8.4 |
| Povidone-Iodine | -3.43 | 3.09 mM | -1.7 |
| Sertaconazole | -7.25 | 4.81 μM | -8.2 |
| Sulconazole | -7.15 | 5.74 μM | -7.7 |
| Terbinafine | -7.26 | 4.75 μM | -6.9 |
| Terconazole | -7.49 | 3.24 μM | -8.8 |
| Tioconazole | -6.78 | 10.66 μM | -6.9 |
| Tolnaftate | -7.25 | 4.82 μM | -7.8 |
| Undecylenic Acid | -5.04 | 202.61 μM | -6.0 |
| Voriconazole | -5.03 | 206.50<br>μM | -6.7 |
| Zinc Undecylenate | 4.54 | - | -1.3 |
| **Antineoplastic drugs** |  |  |  |
| Imatinib | **-9.80** | 65.97 nM | **-9.4** |
| Pazopanib | -9.40 | 128.95<br>nM | -8.7 |
| Regorafenib | -8.56 | 533.39<br>nM | -8.9 |
| Sorafenib | -8.45 | 640.70<br>nM | -8.3 |
| Sunitinib | -7.34 | 4.14 μM | -7.8 |

### 2D Ligand interaction diagram

The antifungal drug Itraconazole with best binding energy and docking position was viewed in a 2D Ligand interaction diagram (Fig. 6). Here, the ligand interactions were presented with conventional hydrogen bonds at A-585 positions of the ARG, Pi-Sigma interactions with two amino acids such as A-586 positions of TRP and A-965 position of ILE, and Alky interactions at A-965 ILE and A: 831 ILE.

**Fig. 6.**
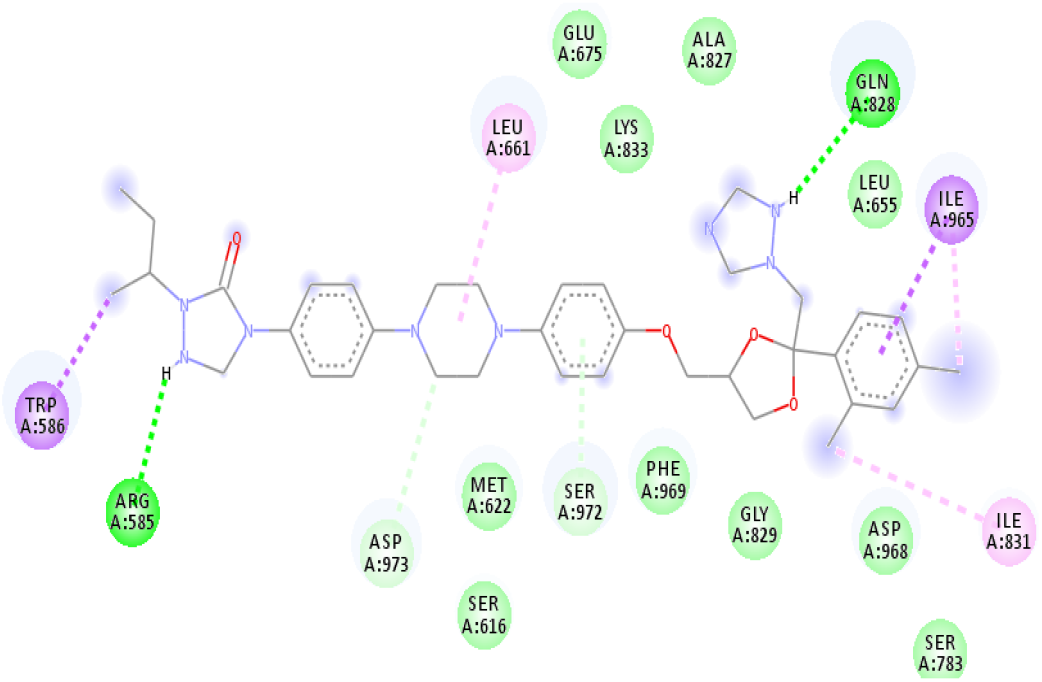
2D diagram of Itraconazole interaction with target protein PDGFRA.

2D Ligand interaction diagrams for docking poses of Imatinib with target protein atoms are shown in Fig. 7. The conventional hydrogen bond at A-623 LYS, unfavourable donor-donor interactions at A-833 position of LYS, Pi-Sulfur interaction at A-973 ASP, Pi-Anion interaction at A-622 MET, and Pi-Alkyl interaction at A-586 position of TRP. Based on the 2D diagram of Itraconazole and Imatinib interaction with PDGFRA protein. The structure of Itraconazole was changed as shown in Fig. 8. It was designated as S1 and its IUPAC name was 1-(butan-2-yl)-4-(4-{[2-(2,4-dichlorophenyl)-2-[(1H-1,2,4-triazol-1-yl)methyl]-1,3-dioxolan-4-yl]methoxy}piperazin-1-yl)-4,5-dihydro-1H-1,2,4-triazol-5-one.

**Fig. 7.**
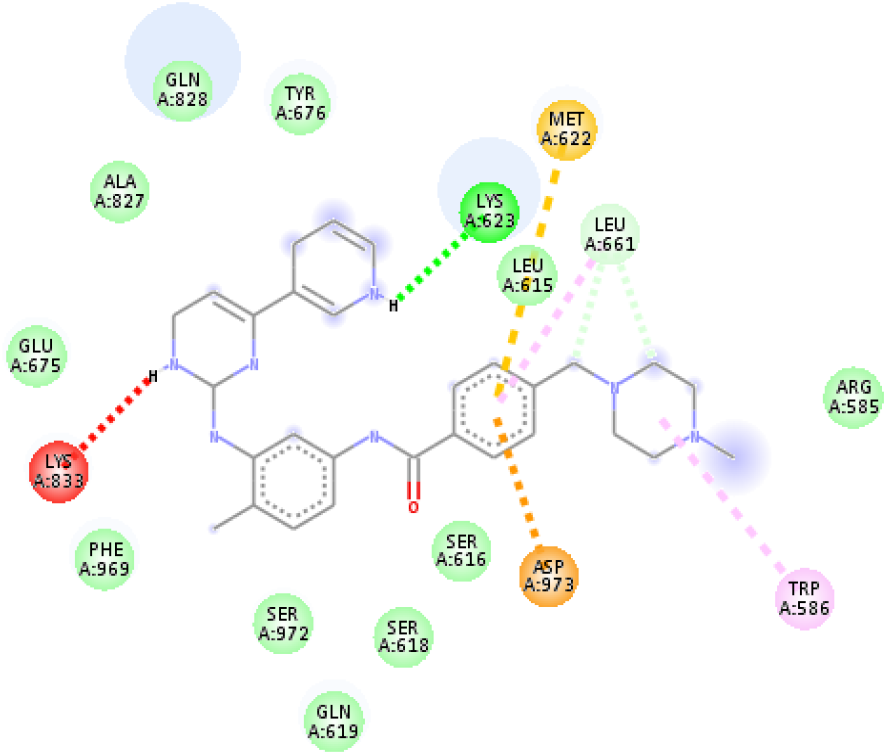
2D diagram of Imatinib interaction with the target protein PDGFRA.

**Fig. 8.**
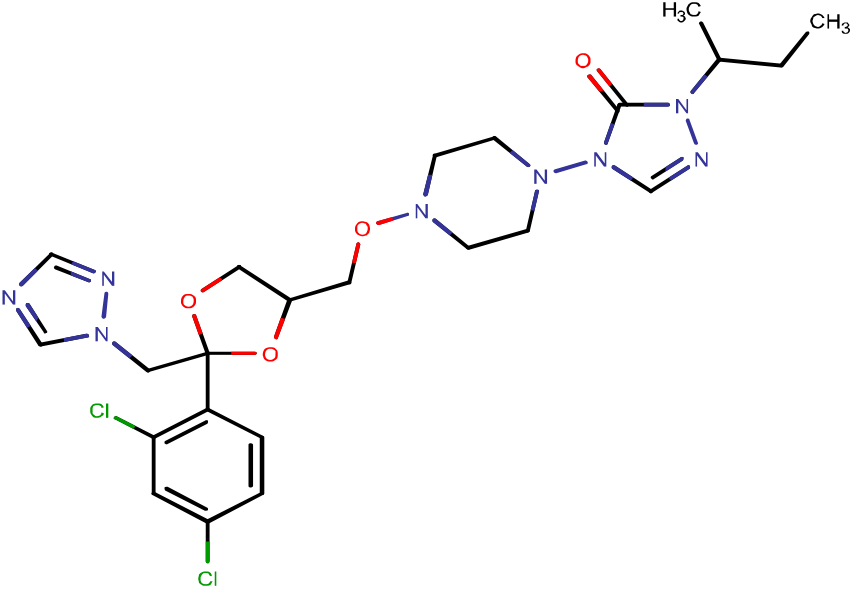
2D structure of S1, Elemental analysis by MarvinSketch-Molecular weight: 553.45, Formula: C_23_H_30_Cl_2_N_8_O_4_, Atom count: 67.

From Imatinib, structure S2 (IUPAC name 1-benzyl-4-methylpiperazine was obtained as shown in Fig. 9.

**Fig. 9.**
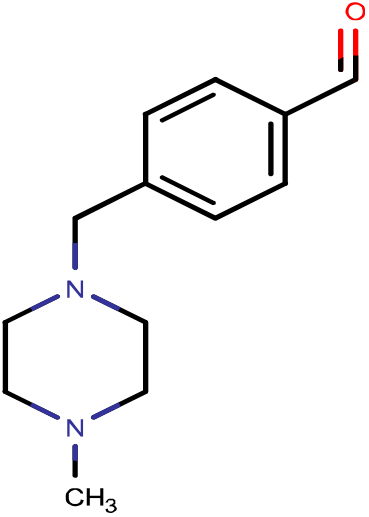
2D structure of altered Imatinib (S2), Elemental analysis by MarvinSketch-Molecular weight: 218.300, Formula: C_13_H_18_N_2_O, Atom count: 34.

Based on the AD and ADV, the docking score for S1 and S2 was not promising. It may be due to their molecular weight and length. So, from S1 and S2 structure, a structure S3 was proposed (Fig. 10). Its IUPAC name was 2-(4-{[2-(2,4-dichlorophenyl)-2-[(1 H-1,2,4-triazol-1-yl)methyl]-1,3-dioxolan-4-yl]methoxy}piperazin - 1 - yl) - 10 - methyl - 8 - [(4 - methylcyclohexyl)methyl] - 1H,2H,3H,5H, 10H-[1,2,4]triazolo[1,2-b]phthalazine-1,5-dione.

**Fig. 10.**
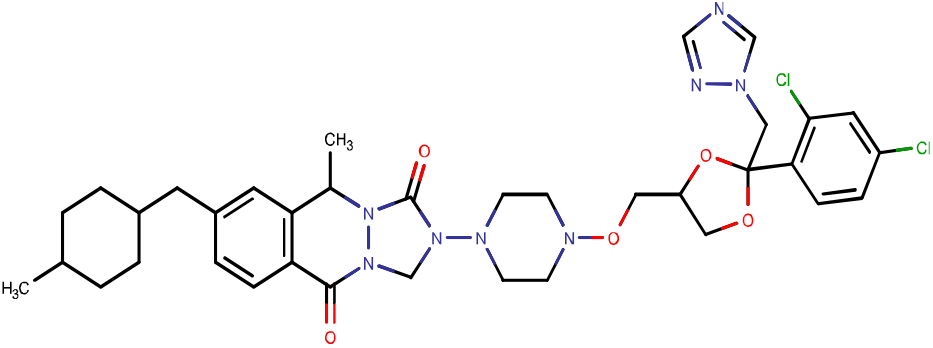
2D structure of S3, Elemental analysis by MarvinSketch - Molecular weight: 739.70, Formula: C_36_H_44_Cl_2_N_8_O_5_, Atom count: 95.

Structure S3 showed −11.47 kcal/mol binding energy and 3.92 nM inhibition constant in AD. Binding energy was −11.0 kcal/mol in ADV docking (Table 2). This predicted value was better than the Imatinib binding energy and inhibition constant for PDGFRA inhibition.

**Table 2.** Binding affinity of proposed ligand and drugs

| Ligand | AutoDock |  | AD Vina |
| --- | --- | --- | --- |
| Antifung | kcal/mo | K<sub>i</sub> | kcal/mol |

| al drugs | I |  |  |
| --- | --- | --- | --- |
| S1 | -7.85 | 1.77 μM | -8.6 |
| S2 | -7.36 | 4.03 μM | -5.8 |
| S3 | -11.47 | 3.92 nM | -11.0 |

The tyrosine kinase inhibitor (TKI) Imatinib interacts with five major CYP isoforms. Use of TKIs with other drugs that decrease absorption or induce metabolism of TKI may result in sub-therapeutic levels of the drug and bring about a decrease in TKI effect. To the contrary, drugs that inhibit the TKI metabolism may cause supra-therapeutic drug levels and toxicity. These interactions possibly lead to a loss of therapeutic effects of TKIs or cause severe to fatal side effects. The suggested antifungal drug-Itraconazole inhibits only two isoforms (see Table 3). Long-term use of Imatinib increases a potential drug-drug interaction risk. Itraconazole, can aid in lowering toxicity rates and providing efficient antitumor therapy.

## DISCUSSION

GISTs are commonly found in adult patients aged between 40 and 70 years with some rare cases of this tumor being developed in children and young adults with similar frequencies in men and women [20]. Studies estimated the rate of GISTs at 7-20 cases per million population years in the United States (Ma et al., 2015). A Swedish study estimated the frequency of GIST at approximately 14.5 cases per million population years, and an Icelandic study reported an incidence of 11 cases per million population years [21][22]. A study conducted in Alberta found the incidence rate of GISTs as 0.91 per 10^5^ population-years [23]. PDGFRA gene mutations that increase the risk of developing GISTs are inherited from a parent, leading to familial GISTs. Through signal transduction, RTK transmits signals from the cell surface into the cell. This activation of the PDGFRA protein then activates several other proteins through phosphorylation, ultimately, triggering a succession of proteins in various signalling pathways.

*In silico* based therapeutic switching of drugs that are already utilized clinically to identify novel therapeutic uses is an upcoming approach as these FDA approved drugs are easily available on the market and have known pharmacokinetic as well as toxicological effects [24]. Therapeutic switching reduces the steps and costs required for drug development and approval processes considerably [25][26]. In this study, the screening of test compounds against PDGFRA showed a best binding affinity for Itraconazole (antifungal) with −9.6 kcal/mol in ADV. Itraconazole was shown to be active against another cancer, the non-small cell lung cancer[27][18]. It was repurposed as an anticancer agent and tested in clinical trials for the treatment of gastric, pancreatic, oesophageal, lung, prostate, and basal cell cancers recently (Tsubamoto et al., 2017). However, there are no studies concerned with GISTs as yet. The binding affinity of the drug Imatinib (antineoplastic) was close to that of Itraconazole. Imatinib, the first targeted agent, represents a promising treatment modality in patients affected by GISTs and chronic myeloid leukemia (CML) [1]. Qiu and his co-workers suggest that genetic polymorphisms may influence the Imatinib-induced ophthalmo-logical toxicities in GIST patients. In addition, Imatinib has several recognized adverse effects, commonly comprising fluid retention with nausea, periorbital edema, abdominal pain, vomiting, muscle cramps, skin rash, headache, diarrhea, and dizziness [28]. The major drawback with Imatinib is resistance development which is therapeutically challenging [29]. The development of specific tyrosine kinase inhibitors, such as Imatinib mesylate, has led to a breakthrough in the treatment of advanced GISTs. Treatment with the drug Imatinib mesylate led to significant improvements in patient survival, with overall response rates in excess of 80 % [30]. Itraconazole as a therapeutic agent against GISTs by inhibiting PDGFRA strongly supports the potential of clinical antineoplastic testing of repurposed drugs such as antifungal drugs.

In conclusion, screening of new effective inhibitors of specific molecular targets in disease treatment can be effectively done by *in silico* approaches. In this study, a suitable system of identifying and screening of PDGFRA inhibitors for the treatment of GISTs by repurposing FDA approved drugs using a molecular docking approach is proposed. Protein stability was checked and key active site residues were identified. Analysis of ligands in the active site aided in the identification of key residues and their interactions for virtual screening. This holds for FDA approved drugs with PDGFRA inhibiting potential. Free binding energy scores based on the docking results showed that among antifungal drugs, Itraconazole had the highest binding energy and stability at the active site. Imatinib exhibited highest binding affinity among the selected antineoplastic drugs for PDGFRA inhibition. These results were validated by correlating the binding affinity from AD and ADV programs used for docking, thereby ensuring that this *in silico* docking based approach is reliable for the suggestion of therapeutic switching. Thus, an as yet antifungal drug, Itraconazole, was shown to be a potentially better PDGFRA inhibitor than Imatinib as the official drug for this purpose. This result suggests Itraconazole for clinical testing in the treatment of GIST. *In silico* analysis also supported the potential of PDGFRA as a promising target for the treatment of GISTs for future *in silico* and *in vitro* strategies for drug design and development.

## Acknowledgment

This work was supported by an International Student fellowship from KMU-Kaohsiung Medical University to the Department of Medicinal and Applied Chemistry.

---

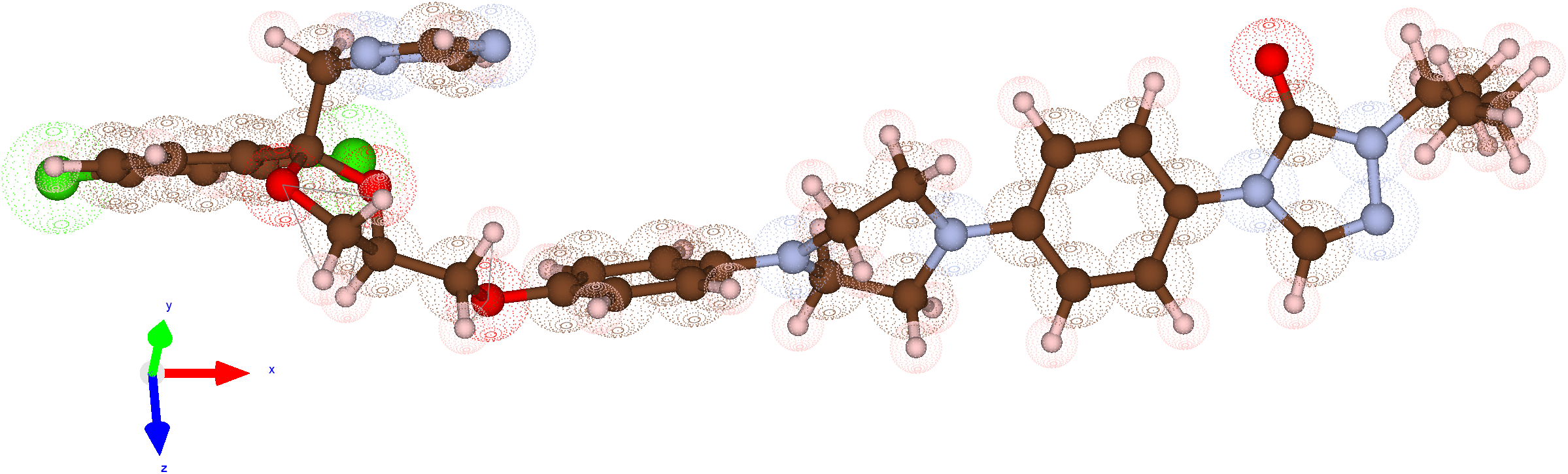

